# Hyaluronidase impacts exposures of long-acting injectable paliperidone palmitate in rodent models

**DOI:** 10.1101/2024.03.03.583160

**Authors:** Henry Pertinez, Amit Kaushik, Paul Curley, Usman Arshad, Eman El-Khateeb, Si-Yang Li, Rokeya Tasneen, Joanne Sharp, Edyta Kijak, Joanne Herriott, Megan Neary, Michaël Noë, Charles Flexner, Eric Nuermberger, Andrew Owen, Nicole C. Ammerman

**Author notes:** **Corresponding Author:** Nicole C. Ammerman.

## Abstract

A significant challenge in the development of long-acting injectable drug formulations, especially for anti-infective agents, is delivering an efficacious dose within a tolerable injection volume. Co-administration of the extracellular matrix-degrading enzyme hyaluronidase can increase maximum tolerable injection volumes but is untested for this benefit with long-acting injectable formulations. One concern is that hyaluronidase could potentially alter the tissue response surrounding an injection depot, a response known to be important for drug release kinetics of long-acting injectable formulations. The objective of this pilot study was to evaluate the impact of co-administration of hyaluronidase on the drug release kinetics, pharmacokinetic profiles, and injection site histopathology of the long-acting injectable paliperidone palmitate for up to four weeks following intramuscular injection in mouse and rat models. In both species, co-administration of hyaluronidase increased paliperidone plasma exposures the first week after injection but did not negate the overall long-acting release nature of the formulation. Hyaluronidase-associated modification of the injection site depot was observed in mice but not in rats. These findings suggest that further investigation of hyaluronidase with long-acting injectable agents is warranted.

## 1. Introduction

Long-acting injectable (LAI) formulations of anti-infective medications hold great promise for simplifying administration and improving adherence, especially for indications in which preventive or therapeutic regimens require prolonged durations (weeks to months) of drug administration, including HIV prevention and treatment [1], malaria prophylaxis [2], viral hepatitis treatment [3, 4], and tuberculosis prevention and treatment [5]. A significant challenge in development of anti-infective LAI formulations is achieving efficacious drug exposure profiles within a tolerable injection volume. A recommended maximum intramuscular (IM) injection volume in adults is 5 mL, although IM injections greater than 3 mL are rare and thus many health care professionals are not familiar with administering larger injection volumes [6]. This is reflected in proposed target product profiles for LAI formulations for malaria and tuberculosis indications, which suggest that, as a minimum requirement, IM injection volume should not exceed 2 mL, and ideally would be <2 mL in adults [5, 7].

One component of the extracellular matrix and interstitial space that contributes to these volume limitations is hyaluronic acid, which is a viscous disaccharide polymer that retains water molecules and prevents sudden bulk fluid flow through the extracellular matrix. This property of hyaluronic acid allows the extracellular matrix to resist compressive forces, such as those created by an LAI drug depot. However, hyaluronic acid can be transiently depolymerized by hyaluronidases, which are enzymes that function naturally as spreading agents to aid in the dispersion of venoms, toxins, and sperm. A recombinant human hyaluronidase (rHuPH20, Hylenex^®^) has been approved by the United States Food and Drug Administration for use as an adjuvant to increase the dispersion and absorption of drugs [8], and the manufacturer of rHuPH20, Halozyme, has developed ENHANZE^®^ drug delivery technology for co-formulation with [non-long-acting] medications to permit significantly larger subcutaneous injection volumes [9]. Therefore, we hypothesized that co-administration of a hyaluronidase could also increase the tolerable IM injection volume for LAI formulations.

The nature of LAI drug formulations is that the drug is slowly released over time from the injected drug depot. Preclinical studies of the LAI antipsychotic agent paliperidone palmitate in a rat model demonstrated that following injection into the thigh muscle, a granuloma forms around the depot site, and co-administration of agents that interfere with the host inflammatory response and abrogate granuloma formation can alter the paliperidone release pharmacokinetics (PK) and lead to lower drug exposures [10, 11]. Although not directly anti-inflammatory, co-administration of hyaluronidase could potentially influence the nature of the immune response following injection of an LAI formulation by affecting immune traffic through the extracellular matrix and interstitial space. The objective of this study was to evaluate the paliperidone release kinetics and PK profile following IM injection with paliperidone palmitate with or without co-administration of hyaluronidase in both mouse and rat models. Paliperidone palmitate was used as a representative approved and marketed LAI formulation for which PK and drug depot histopathology data in rat models were published [10, 11]. In this study, we also included assessment in a mouse model, as mouse models are often used in pre-clinical treatment models of infectious diseases, including mouse models of tuberculosis used to evaluate developmental LAI formulations of anti-tuberculosis drugs [12–14].

## 2. Materials and Methods

### 2.1. Mice (sourcing, husbandry, and other general information)

All studies using mice were conducted at the Johns Hopkins University Center for Tuberculosis Research, and all study procedures were approved by the Institutional Animal Care and Use Committee of the Johns Hopkins University. Adult (6-8 weeks old) female BALB/c mice, purchased from Charles River, were used in all studies. All mg/kg dosing of mice was based on an average mouse body weight of 20 g. Mice were housed in individually ventilated cages with 4 mice (one sampling group) per cage with access to food and water *ad libitum* and with sterile shredded paper for bedding. Room temperature was maintained at 22-24°C, and a 12-hour light/12-hour dark cycle was used. Mice were killed by exsanguination via cardiac puncture under inhalation anesthesia with isoflurane, followed by cervical dislocation.

### 2.2. Rats (sourcing, husbandry, and other general information)

All studies using rats were conducted at the University of Liverpool, under the UK Home Office Project License PP9284915 in accordance with the Animals Scientific Procedures Act (ASPA) 1986 and approved by the local Animal Welfare and Ethical Review Board (AWERB). Adult (6-8 weeks old) male Sprague Dawley rats, purchased from Charles River, were used in all studies. All mg/kg dosing was based on an average body weight of 250 g. Rats were housed in individually ventilated cages with 4 rats per cage and access to food and water *ad libitum*. Sterile wood chippings, nesting material and enrichment was readily available in all cages. Room temperature was maintained at 20-24°C, 45-65% humidity and a 12-hour light/dark cycle was used. Rats were killed by exsanguination via cardiac puncture under inhalational anesthesia, followed by cessation of breathing as confirmation of death.

### 2.3. Sourcing and preparation of paliperidone palmitate and hyaluronidase for IM administration and of paliperidone for IV administration to mice

The paliperidone palmitate LAI formulation, manufactured by Janssen Pharmaceutica NV and sold under the brand name INVEGA SUSTENNA^®^, was supplied at 156 mg/mL in an aqueous IM-injectable suspension containing the following inactive ingredients: polysorbate 20, polyethylene glycol 4000, citric acid monohydrate, sodium dihydrogen phosphate monohydrate, sodium hydroxide, and water for injection. INVEGA SUSTENNA^®^ pre-filled syringes were purchased from the Johns Hopkins Hospital Pharmacy. Allometric scaling based on the standard rat dose of 20 mg/kg [10, 11] and standard first human dose (234 mg or about 4 mg/kg) [15] suggested a dose of 40 mg/kg in mice. However, administration of this dose would require injecting only 5 µL of the INVEGA SUSTENNA^®^ suspension. We were concerned about accuracy in injecting such a small volume into mouse thighs. However, as we did have previous experience with 8 µL IM injections in mice [12, 13], a pragmatic decision to dose with 8 µL of the INVEGA SUSTENNA^®^ suspension was made, resulting in a 62.4 mg/kg dose in the mice.

Hyaluronidase powder was purchased from Millipore Sigma, product number H3506 (hyaluronidase from bovine testes); this specific product has been previously used for IM injection in mice [16]. The powder was dissolved in normal saline at concentrations of 2.5 and 15 enzyme units (U) per microliter, resulting in doses of 5 U and 15 U, respectively, in a 2 µL volume. These doses of hyaluronidase had been previously administered to mice by IM injection [16–18]. The solutions were filter-sterilized before use.

For administration of the paliperidone palmitate alone or with buffer (normal saline with or without hyaluronidase), the INVEGA SUSTENNA^®^ pre-filled syringe was vigorously shaken for 10 seconds to ensure a homogeneous suspension, according to the prescribing information [15]. The suspension was then carefully expelled and aliquotted into four sterile microcentrifuge tubes. In one tube, the suspension was used directly for injection. In the remaining three tubes the paliperidone palmitate suspension was mixed by gentle pipetting with buffer (normal saline) alone or buffer containing hyaluronidase at 2.5 or 15 U/µL, with buffer representing 20% (vol/vol) of the final volume. For IM injection of mice, 8 µL of paliperidone palmitate suspension alone or 10 µL of the paliperidone palmitate suspension plus buffer (with or without hyaluronidase) were injected into the right hind thigh using a BD Veo insulin syringe with a BD Ultra-Fine 3/10 mL 6-mm by 31-gauge needle with a half-unit scale. This syringe/needle combination had been previously used by our group to successfully administer 8 µL IM injection volumes [12, 13].

Paliperidone powder was purchased from Carbosynth (now Biosynth), product number FP2572. For intravenous (IV) administration, the paliperidone powder was dissolved at a concentration of 0.70 mg/mL in 10 mM sodium acetate-acetic acid buffer pH 4.0 [11], resulting in a dose of 3.5 mg/kg in a 100 µL injection volume. The solution was filter-sterilized before use. This dose was selected to maximize the chances of detection while avoiding the maximum IV dose (10 mg/kg) that Janssen reported testing in mouse toxicity studies [19]. A paliperidone IV injection group was included to enable calculation of the absolute bioavailability of paliperidone in mice and confirm any “flip-flop” kinetics [20] associated with slow depot release. Paliperidone was used for IV administration instead of paliperidone palmitate because the palmitate ester is hydrolyzed as part of overall depot release, freeing active paliperidone moiety systematically for pharmacological effect and quantification. Thus, even when it is paliperidone palmitate that is administered by IM injection, the disposition being monitored and quantified is actually that of paliperidone, which therefore must be compared with the IV disposition of the same.

### 2.4. Sourcing and preparation of paliperidone palmitate and hyaluronidase for IM administration to rats

The paliperidone palmitate LAI suspension INVEGA SUSTENNA^®^ was purchased from Royal Liverpool Hospital Pharmacy. Based on a previous publication of this formulation by Janssen researchers [10, 11], a standard dose of 20 mg/kg was administered to the rats. Hyaluronidase powder was purchased from Sigma. The powder was dissolved in phosphate buffered saline to generate doses of 5 U and 15 U of hyaluronidase per 2 μL. Rats were administered a 32 μL or 34 μL volume of INVEGA SUSTENNA^®^ or INVEGA SUSTENNA^®^ with hyaluronidase respectively, via IM injection.

### 2.5. Initial toxicity studies in mice

Prior to conducting the PK study in mice, we first assessed the overt tolerability/toxicity in mice following administration of paliperidone palmitate or paliperidone by IM or IV injection, respectively. A total of 10 mice were used for these studies. Five mice were administered paliperidone palmitate alone at 62.4 mg/kg by IM injection. Following injection, the mice were observed every 1-2 hours for the first 8 hours and then daily for up to two weeks, at which point the study was terminated. Five mice were administered paliperidone at 3.5 mg/kg by IV injection. Following injection, the mice were observed every 1-2 hours for the first 8 hours and then checked again at 24 hours, at which point the study was terminated.

### 2.6. PK study in mice

The primary study outcome was plasma paliperidone concentrations. Mice (n = 64) were randomly assigned to one of the following five groups: (1) paliperidone palmitate alone at 62.4 mg/kg by IM injection (12 mice); (2) paliperidone palmitate 62.4 mg/kg plus buffer without hyaluronidase by IM injection (12 mice); (3) paliperidone palmitate 62.4 mg/kg plus 5 U hyaluronidase by IM injection (12 mice); (4) paliperidone palmitate 62.4 mg/kg plus 15 U hyaluronidase by IM injection (12 mice); (5) paliperidone solution at 3.5 mg/kg by IV injection (16 mice). For mice receiving paliperidone palmitate alone or with buffer/hyaluronidase by IM injection, blood samples were obtained from mice at 0.25, 1, 2, 4, 7, 10, 24, 48, 72, 96, 168, 336, and 672 hours post injection. For mice receiving paliperidone by IV injection, blood samples were obtained from mice at 0.1, 0.25, 0.5, 1, 2, 4, 7, 10, and 24 hours post injection. For each group at each time point, blood was sampled from four mice via in-life mandibular bleeding, except for the final time point, when blood was drawn by cardiac puncture. The study design, including how the mice were serially sampled and at what time point each group of mice received cardiac punctures, is detailed in **Supplemental Table S1**. Blood was collected into BD vacutainer plasma separation tubes with lithium heparin, and plasma was separated by centrifugation at 15,000 × *g* for 10 min at room temperature. Plasma was transferred to 1.5-mL O-ring screw-cap tubes and stored at −80°C. Frozen samples were shipped on dry ice to the University of Liverpool for processing and determination of paliperidone concentration.

### 2.7. PK study in rats

Because there are multiple published reports for IM administration of paliperidone palmitate and IV administration of paliperidone in rats [10, 11, 21], initial tolerability studies in rats were not deemed necessary in this project. The primary study outcome was plasma paliperidone concentrations. Rats (n = 12) were randomly assigned to one of the following three groups: (1) paliperidone palmitate alone at 20 mg/kg by IM injection (4 rats); (2) paliperidone palmitate 20 mg/kg plus 5 U hyaluronidase by IM injection (4 rats); (3) paliperidone palmitate 20 mg/kg plus 15 U hyaluronidase by IM injection (4 rats). Serial blood samples were obtained from each rat at 0.25, 1, 2, 4, 7, 10, 24, 48, 72, 96, 168, 360, 504, and 672 hours post injection. For each group at each time point, blood was sampled via lateral tail vein bleed, except for the final time point, when blood was drawn by cardiac puncture. The study design, including how the rats were serially sampled, is detailed in **Supplemental Table S2**. Blood was collected into 1 mL syringes via a 25-gauge needle and transferred to an Eppendorf tube containing heparin (Starstedt, UK). Plasma was separated by centrifugation at 13,000 × *g* for 3 min at room temperature. Plasma was transferred to 1.5-mL Eppendorf tubes and stored at −80°C until analysis.

### 2.8. Determination of paliperidone plasma concentrations

The plasma concentration of paliperidone in both rats and mice was quantified using a liquid chromatography tandem mass spectrometric method, which conforms to United States Food and Drug Administration bioanalytical development guidelines [22]. A stock solution of 1 mg/mL paliperidone was prepared in methanol and stored at 4°C until used for preparation of a standard curve from 500 ng/mL to 1.9 ng/mL in either rat or mouse plasma by serial dilution. Paliperidone was extracted from blood plasma samples via protein precipitation: in glass tubes, 100 μL of sample (blank matrix, calibrator standard, quality control, or unknown) was combined with 450 μL acetonitrile and 50 μL internal standard stock solution (50 ng/mL sofosbuvir). Following centrifugation at 3,000 × *g* for 5 minutes, 80 μL of the supernatant fraction and 20 μL H_2_O were placed in chromacol vials for analysis. Quantification was then performed using a SCIEX 6500+ QTRAP (SCIEX MA, USA) operating in positive mode (m/z 427 > 207 and 110).

Chromatographic separation was achieved using a multi-step gradient with a Kinetex 2.6 µm C18 column (Phenomenex, Cheshire, UK) using mobile phases A (100% H_2_O, 0.05% formic acid) and B (100% acetonitrile, 0.05% formic acid). Chromatography was conducted over 3.5 minutes at a flow rate of 400 μL/min. Concentrations obtained were expressed in ng/mL.

### 2.9. PK analyses

Plasma concentration-time profiles were analyzed with non-compartmental PK analysis in the R data analysis environment [23] and Microsoft Excel for calculation of terminal half-lives (T_1/2_) and other PK parameters including maximum plasma concentration (C_max_) and its time of occurrence (T_max_) and area under the time-concentration curve for partial sections of the time course profile (AUC_0-t_), the full profile (AUC_0-last_) and the full profile extrapolated to infinity (AUC_inf_). For mouse data, where a maximum of three plasma samples were taken from any individual mouse over the course of the study, the composite mean PK profile from all mice was analyzed and PK parameters calculated. For rat data where full time courses could be sampled in each individual animal, mean PK parameter values and their standard deviations were calculated following from analysis of individual PK profiles. For calculation of bioavailability (F) the contribution of the palmitate ester to the molecular weight of paliperidone palmitate (such that paliperidone palmitate is 64.1% paliperidone by mass) was accounted for in dose normalization such that 62.4 and 20 mg/kg doses of paliperidone palmitate are equivalent to 40 and 12.8 mg/kg doses of paliperidone, respectively. Plasma concentrations following IV injection of paliperidone in rat and calculation of bioavailability was sourced from the published data of Sun *et al*. [21].

### 2.10. Histopathology assessment

A secondary study outcome was a descriptive assessment of the histopathology surrounding the drug depot following intramuscular injection. For both mice and rats, the hind thighs that received IM injections were removed following cardiac puncture and confirmation of death and fixed in formalin. Fixed rat tissues were shipped to Johns Hopkins University for analysis. Fixed mouse and rat thigh samples were leveled at 60 µm and sectioned at a thickness of 5-6 µm for hematoxylin and eosin staining. Sections were processed, stained, and scanned by the Johns Hopkins University Oncology Tissue Services.

## 3. Results and discussion

### 3.1. Initial toxicity studies of paliperidone palmitate (IM) and paliperidone (IV) in mice

Because the proposed 62.4 mg/kg IM dose of paliperidone palmitate was based on allometric scaling while using a practical injection volume, we did not know if this dose would be tolerated by the mice. Therefore, prior to conducting the full PK study, we first assessed the overt toxicity in mice following IM injection of paliperidone palmitate at 62.4 mg/kg. None of the five mice injected exhibited any signs of toxicity at any time during the two-week follow-up period post injection. Although our proposed dose of 3.5 mg/kg of paliperidone in solution for IV injection was selected to stay below the maximum IV dose (10 mg/kg) reported to have been evaluated in mouse toxicity studies according to a regulatory report [19], we also wanted to first assess the tolerability of this dose in mice before initiating our full PK study. Immediately following IV injection, four out of the five mice became lethargic and exhibited ptosis, which was consistent with the toxicity information in the regulatory report. By five hours post injection, the symptoms had noticeably improved, and by 24 hours post injection, the mice were fully recovered. Based on these observations, we determined that these IM and IV doses of paliperidone palmitate suspension and paliperidone solution, respectively, were suitable without an overt toxicity issue.

### 3.2. Evaluation of the impact of hyaluronidase on paliperidone plasma exposure profiles following IM injection of paliperidone palmitate LAI formulation in mice

Visual inspection of the mouse plasma PK profile time courses indicated a multiphasic release profile for paliperidone following IM dosing as paliperidone palmitate in mice (**Fig. 1A**), with at least 2 or 3 release fraction processes leading to local maxima (peaks) in the concentration-time profile at time points as follows: paliperidone palmitate alone: 7 h, 1 day and 1 week; paliperidone palmitate + buffer: 2 h, 2 days and 2 weeks; paliperidone palmitate + 5 U hyaluronidase: 2 h, and 1 week; paliperidone palmitate + 15 U hyaluronidase, 10 h and 1 week. Because of the limitations of sparse sampling, the full shape of the complex PK profile and the true C_max_ may not have been observed with the time points taken. All individual mouse data are available in **Supplementary Data File S1**.

**Figure 1.**
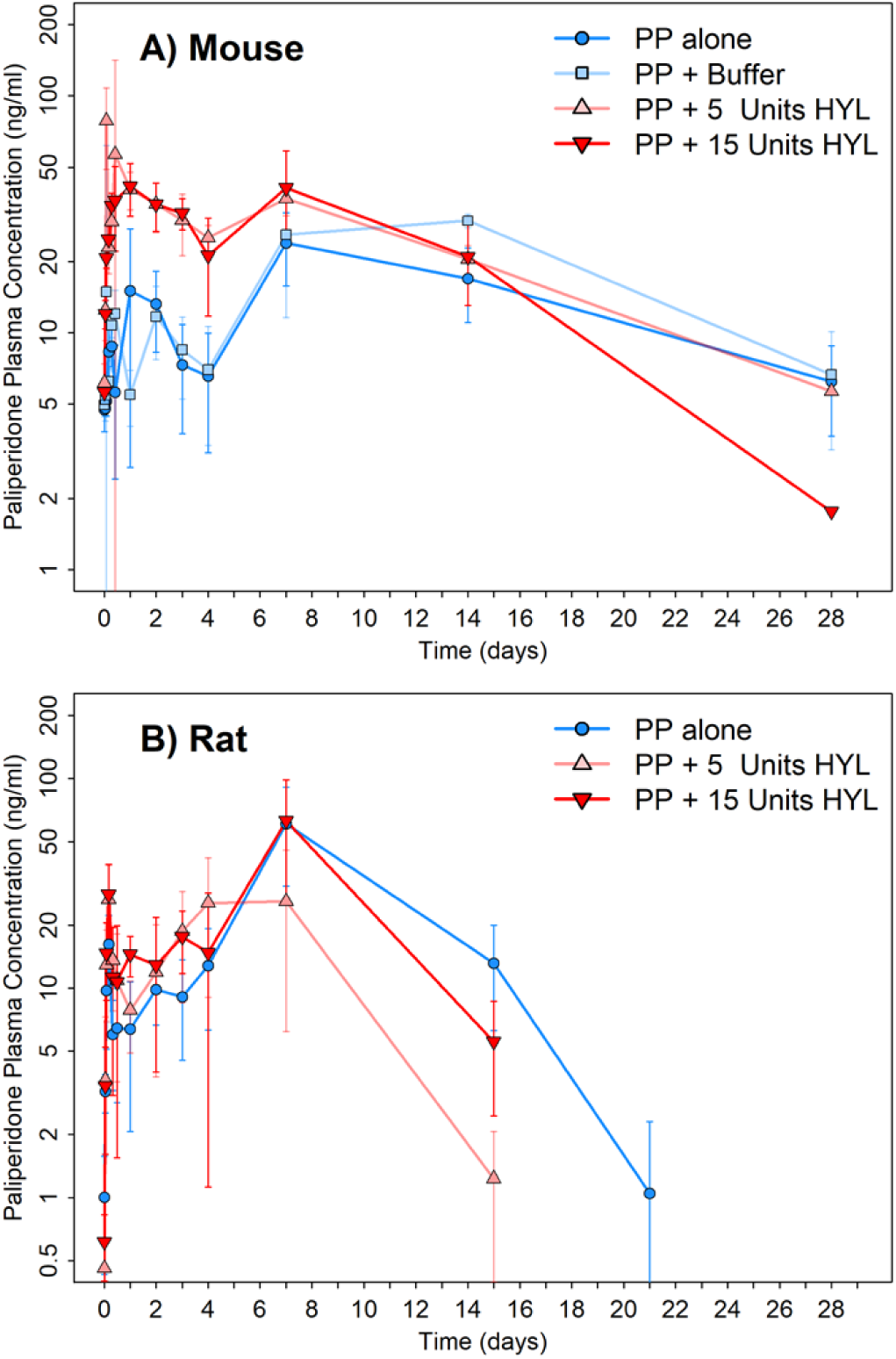
Plasma concentration-time PK profiles for paliperidone in A) mice and B) rats following IM injection with paliperidone palmitate (PP) with or without hyaluronidase (HYL). Data points represent mean values (n = 4 samples per time point for mice; n = 4 samples per time point for rats), and error bars represent +/- standard deviation. All individual mouse and rat data are available in **Supplementary Data Files S1 and S2**, respectively.

Co-administration of hyaluronidase increased exposure in the first 4 days of the time course with AUC_0-4d_ values approximately 3.5 to 4x greater in groups receiving paliperidone palmitate with hyaluronidase compared to the group receiving paliperidone palmitate without hyaluronidase (**Table 1**). A similar effect was observed with either 5 or 15 U hyaluronidase. Although there were multiple local concentration in the profiles, the overall C_max_ was also approximately 2 to 3x greater when hyaluronidase was co-administered (**Table 1**; **Fig. 1A**). Co-administration of hyaluronidase did not compromise the long-acting exposures from IM administration of this formulation to mice, as prolonged exposures were still maintained for 28 days, with similar AUCs, bioavailability (F ∼30%.), and terminal half-lives of (100-250 h) in the presence or absence of hyaluronidase (**Table 1**). In all cases the terminal half-lives of paliperidone following IM dosing as the palmitate ester were confirmed as “flip-flop” [20] in nature when taken in comparison to the IV disposition half-life of paliperidone in mice of 4.5 h (**Table 2**; **Supplementary Fig. S1**).

**Table 1.** PK parameters for paliperidone following IM injection as paliperidone palmitate with or without hyaluronidase in mice and rats. For the mouse data, the composite mean profile parameters (n=12 mice) are presented. For the rat data, the mean and [standard deviation] from n = 4 individual rats are provided for each parameter.

| Species | Study group | AUC<sub>0-96</sub><br>(ng·h/mL) | AUC<sub>0-last</sub><br>(ng·h/mL) | AUC<sub>0-inf</sub><br>(ng·h/mL) | F<br>(%) | Terminal<br>T<sub>1/2</sub><br>(h) | C<sub>max</sub><br>(ng/mL) | T<sub>max</sub><br>(h) |
| --- | --- | --- | --- | --- | --- | --- | --- | --- |
| Mouse | 1) IM paliperidone palmitate 62.4 mg/kg alone | 967 | 9391 | 11693 | 25 | 256 | 23.9 | 168 |
|  | 2) IM paliperidone palmitate 62.4 mg/kg + saline buffer | 851 | 12820 | 14316 | 30 | 156 | 29.7 | 336 |
|  | 3) IM paliperidone palmitate 62.4 mg/kg + 5 U hyaluronidase | 3720 | 15143 | 16661 | 35 | 186 | 96.5 | 10 |
|  | 4) IM paliperidone palmitate 62.4 mg/kg + 15 U hyaluronidase | 3165 | 14396 | 14671 | 31 | 108 | 41.5 | 24 |
| Rat | 1) IM paliperidone palmitate 20 mg/kg alone | 864<br>[292] | 11679<br>[5109] | 12412<br>[5130] | 52 | 78<br>[27] | 60.8<br>[30] | 168<br>[0] |
|  | 2) IM paliperidone palmitate 20 mg/kg + 5 U hyaluronidase | 1425<br>[711] | 5763<br>[3423] | 7289<br>[2344] | 31 | 48<br>[10] | 33.6<br>[16] | 86<br>[95] |
|  | 3) IM Paliperidone Palmitate 20 mg/kg + 15 U hyaluronidase | 1516<br>[307] | 11285<br>[4633] | 11879<br>[4210] | 50 | 63<br>[31] | 50.3<br>[39] | 127<br>[82] |

**Table 2.**
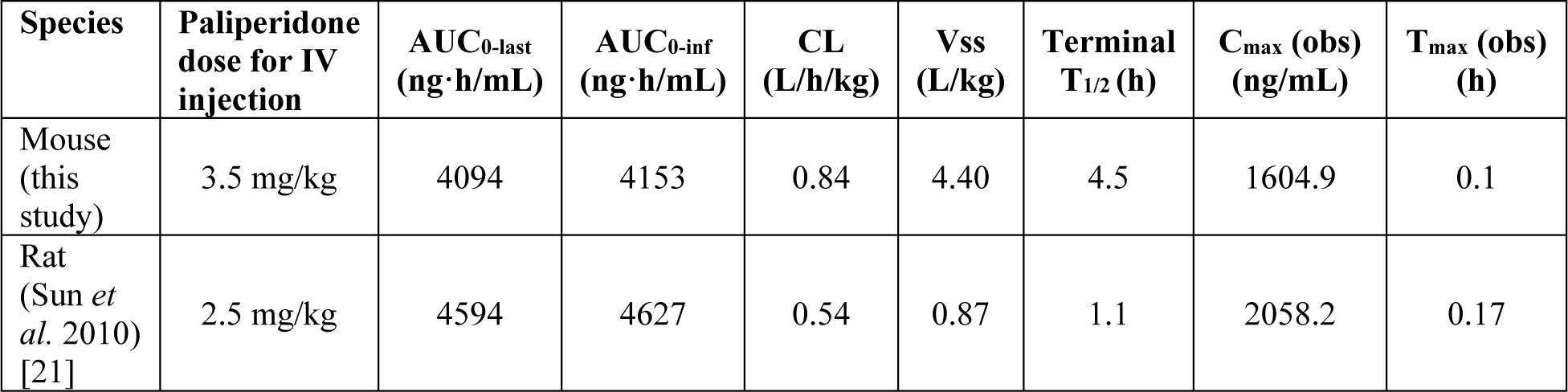
PK parameters for paliperidone following IV doses in mice and rats.

| Species | Paliperidone dose for IV injection | AUC<sub>0-last</sub><br>(ng·h/mL) | AUC<sub>0-inf</sub><br>(ng·h/mL) | CL<br>(L/h/kg) | V<sub>ss</sub><br>(L/kg) | Terminal<br>T<sub>1/2</sub> (h) | C<sub>max</sub> (obs)<br>(ng/mL) | T<sub>max</sub> (obs)<br>(h) |
| --- | --- | --- | --- | --- | --- | --- | --- | --- |
| Mouse<br>(this study) | 3.5 mg/kg | 4094 | 4153 | 0.84 | 4.40 | 4.5 | 1604.9 | 0.1 |
| Rat<br>(Sun *et al.* 2010)<br>[21] | 2.5 mg/kg | 4594 | 4627 | 0.54 | 0.87 | 1.1 | 2058.2 | 0.17 |

### 3.3. Evaluation of the impact of hyaluronidase on paliperidone plasma exposure profiles following IM injection of paliperidone palmitate LAI formulation in rats

Because the IM dose of paliperidone palmitate suspension had been used in previously published studies with Janssen researchers, it was not necessary to conduct an initial toxicity study in rats. As a PK profile following IV injection with paliperidone solution was also already published [21], it was also not necessary to include such a group in this study. To further reduce the number of rats required for this arm of the study, we omitted the paliperidone palmitate plus buffer group, as data from the mouse PK study indicated that adding buffer alone (without hyaluronidase) did not impact outcome, and furthermore the concentration of buffer used with hyaluronidase was proportionally much smaller in the preparations for injections in rats compared to the preparations for injections in mice. All individual rat data are available in **Supplementary Data File S2**.

Similarly for mice, visual inspection of the rat plasma PK profile time courses also indicated a multiphasic release profile for paliperidone following IM dose as paliperidone palmitate, with two apparent release fraction processes leading to local maxima (peaks) at 4h and 1 week in mean concentration-time profiles for all three study groups, with or without co-administration of hyaluronidase (**Fig. 1B**). Also as in mice, co-administration of hyaluronidase increased exposure in the first 4 days of the time course but with reduced apparent effect compared to mice, with AUC_0-4d_ ∼1.6x greater with hyaluronidase present (**Table 1**) with similar effect for either 5 or 15 U hyaluronidase. The reduced apparent effect of hyaluronidase may likely be attributed to the rats receiving a lower weight normalized dose of hyaluronidase (*i.e.*, 5 or 15 U per 0.25 kg typical body weight in rats vs per 0.02 kg typical body weight in mice). Inter-individual variability across animals obscured somewhat the potential effect on overall C_max_ in rats, with outlier individuals with low exposure and truncated profiles due to sampling difficulties (**Supplementary Data File S2**). Similarly, as in mice, limitations of sparse sampling in the study design for welfare/blood volume sampling restriction (and logistical) reasons, may also have meant that the full shape of the complex profile and the true C_max_ may not have been observed with the time points taken.

Co-administration of hyaluronidase again did not appear to compromise the long-acting exposures from IM administration of this formulation to rats, and prolonged exposure was still maintained for ∼20 days (noting the rats received a lower weight normalized dose compared to mice), with similar AUCs, % F (of ∼30-50%.), and terminal half-lives of (50-75 h) in the presence or absence of hyaluronidase. As for mice, the terminal half-lives of paliperidone following IM dosing as the palmitate ester in rat were confirmed as “flip-flop” [20] in nature when taken in comparison to the IV disposition half-life of paliperidone in rats of 1.1 h based on the PK data of Sun *et al*. [21] (**Table 2**).

### 3.4. Histopathological assessment

A secondary outcome in this study was assessment of the histopathology surrounding the injection site. Because a limited number of animals were used in this study (as serial, in-life blood sampling used for assessment of the primary outcome) and also because animals were only sacrificed at the final PK time points, sample numbers for histopathological assessment were low and thus only used for a descriptive, non-quantitative assessment. Additionally, histopathological assessment was not always able to locate the injection site. However, assessment was possible for at least one animal in each study arm. In animals where a clear injection site was found, there was variability in the location of the depot: sometimes the medication was located in the muscle, between the myocytes of the thigh muscles, while in other animals, the medication was found against the muscle, in the nearby adipose tissue (see example in **Supplementary Fig. S2**). It is not clear from pathological assessment whether this variation originates from variation in the injection site, or whether the medication might have been displaced due to the higher pressure in the muscle compared to the adipose tissue (*i.e*., during contraction of the muscle).

In the mouse thighs, it appeared that macrophages had phagocytized almost all the injected material after 168 hours, as seen by the foamy aspect of the cytoplasm of the macrophages (**Fig. 2A-B**; **Fig. S2**). There was no central cavitation, and a very limited amount of other inflammatory cells were present. In mice injected with paliperidone palmitate without hyaluronidase (*i.e*., paliperidone palmitate alone or with buffer only), the collections of foamy macrophages formed a mass with a pushing, smooth border, and very little infiltration between the surrounding individual myocytes (**Fig. 2A**). However, when paliperidone palmitate was co-administered with hyaluronidase, the macrophages showed infiltration between individual myocytes (**Fig. 2B**), suggesting a possible role for hyaluronidase to digest the extracellular matrix and allow the medication to disperse between individual myocytes. A dose-effect with hyaluronidase was not detectable in this small sample set.

**Figure 2.**
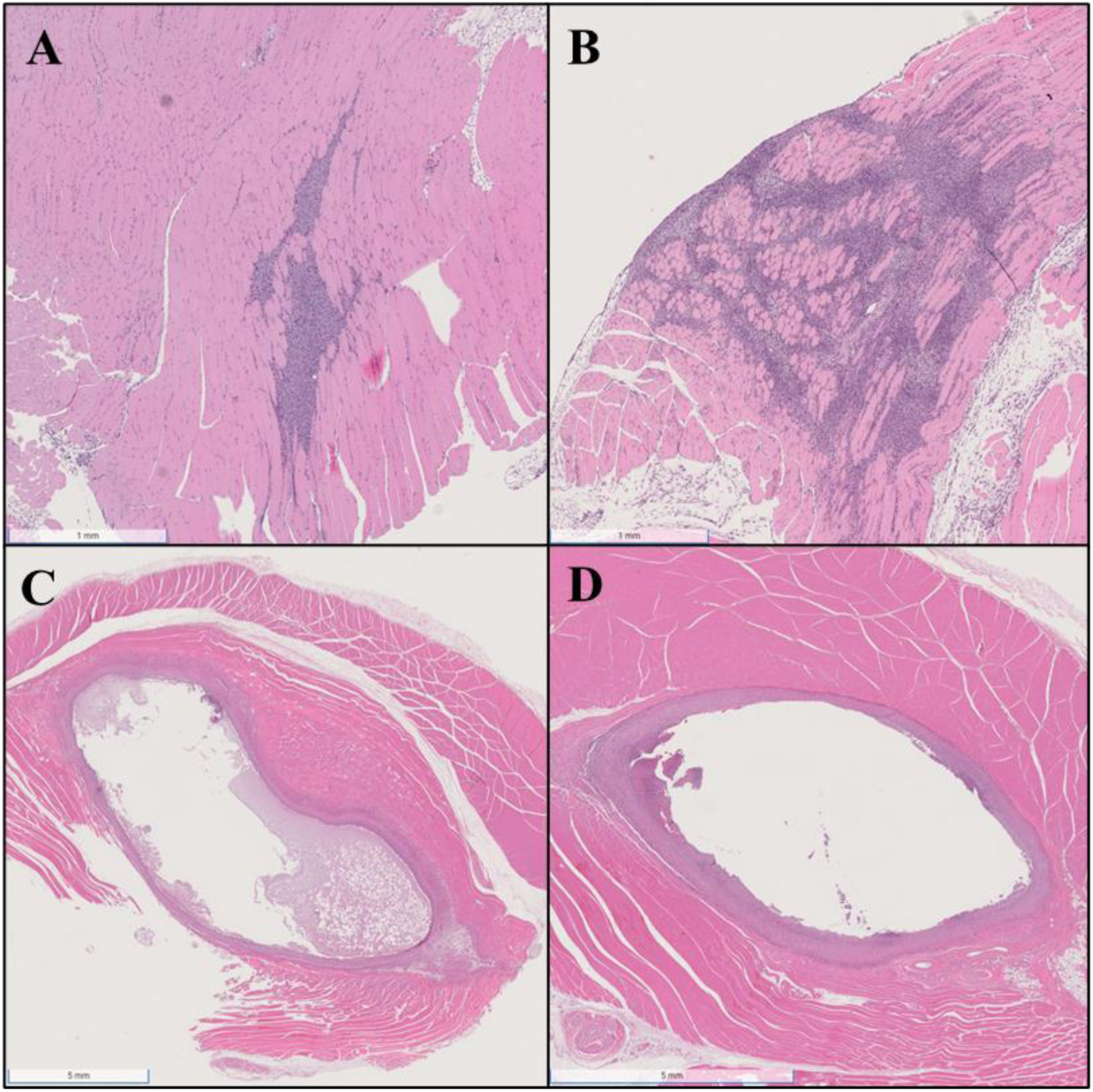
Representative hematoxylin and eosin stained sections showing the paliperidone palmitate depot site in the thigh muscles of mice (A-B) and rats (C-D). Panel A shows the depot site in a mouse thigh that was injected with paliperidone palmitate with buffer only (no hyaluronidase), 336 hours (2 weeks) post injection. The depot is located in macrophages, which form a mass with little infiltration between surrounding myocytes. Panel B shows a mouse thigh that was injected with paliperidone palmitate plus 15 U hyaluronidase, 168 hours (1 week) post injection. The depot is located in macrophages, which show infiltration between surrounding myocytes. Panel C shows a rat thigh that was injected with paliperidone palmitate alone (no buffer, no hyaluronidase), 672 hours (4 weeks) post injection. The depot is surrounded by a wall of macrophages and fibroblasts, forming a cavitating granuloma. No infiltrations are seen between surrounding cells. Panel D shows a rat thigh that was injected with paliperidone palmitate plus 5 U hyaluronidase, 672 hours (4 weeks) post injection. The depot is surrounded by a wall of macrophages and fibroblasts, forming a cavitating granuloma. No infiltrations are seen between surrounding cells.

There was a clear difference between mice and rats regarding the immune reaction to the injection of paliperidone palmitate. In rats, granulomas formed with a central cavity and necrosis, surrounded by a wall of macrophages and fibroblasts, and the cavitating granulomas had smooth borders and showed little infiltration between the individual myocytes of the thigh muscle (**Fig. 2C-D**). This observed granuloma structure aligns well with that previously described by Darville *et al.* following intramuscular injection of paliperidone palmitate in rats [11]. Also in contrast to the mice, co-administration of hyaluronidase did not discernably affect the granulomas or the surrounding myocytes (**Fig. 2C** versus **Fig. 2D**). This apparent lack of impact by hyaluronidase could be due to the hyaluronidase dosing in rats, which, as already discussed, received a lower weight normalized dose than what was used in the mice. This aligns with the reduced impact of hyaluronidase that was observed in the PK results (**Fig. 1**).

## 4. Conclusions

Co-administration of hyaluronidase altered depot release of paliperidone in the first four days (96 h) following IM injection to BALB/c mice and Sprague-Dawley rats, increasing plasma exposures. The higher exposures early in the profile may necessitate further evaluation of the safety and tolerability, however, these data support further investigation of hyaluronidase as an approach to increase injection volumes of other LAIs, particularly those of interest in the anti-infectives therapeutic area such as cabotegravir and rilpivirine.

Multiphasic release processes suggested by the data, with potential modulation by hyaluronidase, are consistent with potential mechanistic descriptions of depot release processes involving resident phagocytic immune cells [11, 24], but this remains to be confirmed.

## CRediT Author Statement

**Henry Pertinez:** Formal analysis, Data curation, Writing - original draft, Writing - review & edit, Visualization

**Amit Kaushik:** Methodology, Investigation, Writing - review & edit.

**Paul Curley:** Investigation, Writing - original draft, Writing - review & edit

**Usman Arshad:** Investigation, Writing - original draft

**Eman El-Khateeb:** Formal analysis, Writing - review & edit

**Si-Yang Li:** Methodology, Investigation, Writing - review & edit.

**Rokeya Tasneen:** Methodology, Investigation, Writing - review & edit.

**Joanne Sharp:** Methodology, Investigation, Data curation, Writing - original draft, Writing - review & edit, Supervision, Project Administration,

**Edyta Kijak:** Investigation, Writing - original draft **Joanne Herriott:** Investigation, Writing - original draft **Megan Neary:** Investigation, Writing - review & edit,

**Michaël Noë:** Investigation, Writing - original draft, Writing - review & edit, Visualization. **Charles Flexner:** Conceptualization, Writing - review & edit, Supervision, Funding acquisition. **Eric Nuermberger:** Conceptualization, Methodology, Resources, Writing - review & edit, Supervision, Funding acquisition.

**Andrew Owen:** Conceptualization, Resources, Writing - original draft, Writing - review & edit, Supervision, Funding acquisition.

**Nicole C. Ammerman:** Conceptualization, Methodology, Data curation, Writing - original draft, Writing - review & edit, Visualization, Supervision, Project Administration, Funding acquisition.

## Acknowledgements

This publication resulted from research supported by the Johns Hopkins University Center for AIDS Research, an NIH funded program (1P30AI094189), which is supported by the following NIH Co-Funding and Participating Institutes and Centers: NIAID, NCI, NICHD, NHLBI, NIDA, NIA, NIGMS, NIDDK, NIMHD. The content is solely the responsibility of the authors and does not necessarily represent the official views of the NIH. The funding institutions had no role in the study design; in the collection, analysis and interpretation of data; in the writing of the report; or in the decision to submit the article for publication.

## Notes

### Competing Interest Statement

The authors have declared no competing interest.

## References

[1] V. Arya, A.C. Hodowanec, S.B. Troy, K.A. Struble, Long-acting formulations for the prevention and treatment of human immunodeficiency virus (HIV)-1 infection: Strategic leveraging and integration of multidisciplinary knowledge to advance public health, Clin Infect Dis, 75 (2022) S498–S501.

[2] J.J. Moehrle, Development of new strategies for malaria chemoprophylaxis: From monoclonal antibodies to long-acting injectable drugs, Trop Med Infect Dis, 7 (2022) 58.

[3] D.L. Thomas, J.J. Kiser, M.M. Baum, Long-acting treatments for hepatitis B, Clin Infect Dis, 75 (2022) S517–S524.

[4] D.L. Thomas, A. Owen, J.J. Kiser, Prospects for long-acting treatments for hepatitis C, Clin Infect Dis, 75 (2022) S525–S529.

[5] N.C. Ammerman, E.L. Nuermberger, A. Owen, S.P. Rannard, C.F. Meyers, S. Swindells, Potential impact of long-acting products on the control of tuberculosis: Preclinical advancements and translational tools in preventive treatment, Clin Infect Dis, 75 (2022) S510–S516.

[6] U. Hopkins, C.Y. Arias, Large-volume IM injections: A review of best practices, Oncol Nurse Advis, Jan-Feb (2013) 32–37.

[7] F. Macintyre, H. Ramachandruni, J.N. Burrows, R. Holm, A. Thomas, J.J. Mohrle, S. Duparc, R. Hooft van Huijsduijnen, B. Greenwood, W.E. Gutteridge, T.N.C. Wells, W. Kaszubska, Injectable anti-malarials revisited: discovery and development of new agents to protect against malaria, Malar J, 17 (2018) 402.

[8] Halozyme, INc. HYLENEX recombinant (hyaluronidase human injection) full prescribing information, 2012. Freely available: https://www.accessdata.fda.gov/drugsatfda_docs/label/2012/021859s009lbl.pdf.

[9] K.W. Locke, D.C. Maneval, M.J. LaBarre, ENHANZE® drug delivery technology: a novel approach to subcutaneous administration using recombinant human hyaluronidase PH20, Drug Deliv, 26 (2019) 98–106.

[10] N. Darville, M. van Heerden, A. Vynckier, M. De Meulder, P. Sterkens, P. Annaert, G. Van den Mooter, Intramuscular administration of paliperidone palmitate extended-release injectable microsuspension induces a subclinical inflammatory reaction modulating the pharmacokinetics in rats, J Pharm Sci, 103 (2014) 2072–2087.

[11] N. Darville, M. van Heerden, D. Marien, M. De Meulder, S. Rossenu, A. Vermeulen, A. Vynckier, S. De Jonghe, P. Sterkens, P. Annaert, G. Van den Mooter, The effect of macrophage and angiogenesis inhibition on the drug release and absorption from an intramuscular sustained-release paliperidone palmitate suspension, J Control Release, 230 (2016) 95–108.

[12] A. Kaushik, N.C. Ammerman, R. Tasneen, S. Lachau-Durand, K. Andries, E. Nuermberger, Efficacy of long-acting bedaquiline regimens in a mouse model of tuberculosis preventive therapy, Am J Respir Crit Care Med, 205 (2022) 570–579.

[13] A. Kaushik, N.C. Ammerman, S. Tyagi, V. Saini, I. Vervoort, S. Lachau-Durand, E. Nuermberger, K. Andries, Activity of a long-acting injectable bedaquiline formulation in a paucibacillary mouse model of latent tuberculosis infection, Antimicrob Agents Chemother, 63 (2019) e00007–00019.

[14] M. Kim, C.E. Johnson, A.A. Schmalstig, A. Annis, S.E. Wessel, B. Van Horn, A. Schauer, A.A. Exner, J.E. Stout, A. Wahl, M. Braunstein, J. Victor Garcia, M. Kovarova, A long-acting formulation of rifabutin is effective for prevention and treatment of *Mycobacterium tuberculosis*, Nat Commun, 13 (2022) 4455.

[15] Janssen Pharmaceutical Companies, INVEGA SUSTENNA® (paliperidone palmitate) extended-release injectable suspension, for intramuscuar use full prescribing information, 2019. Freely available: https://www.accessdata.fda.gov/drugsatfda_docs/label/2017/022264s023lbl.pdf.

[16] P. Hojman, C. Brolin, N. Norgaard-Christensen, C. Dethlefsen, B. Lauenborg, C.K. Olsen, M.M. Abom, T. Krag, J. Gehl, B.K. Pedersen, IL-6 release from muscles during exercise is stimulated by lactate-dependent protease activity, Am J Physiol Endocrinol Metab, 316 (2019) E940–E947.

[17] J.D. Schertzer, D.R. Plant, G.S. Lynch, Optimizing plasmid-based gene transfer for investigating skeletal muscle structure and function, Mol Ther, 13 (2006) 795–803.

[18] J.M. McMahon, E. Signori, K.E. Wells, V.M. Fazio, D.J. Wells, Optimisation of electrotransfer of plasmid into skeletal muscle by pretreatment with hyaluronidase -- increased expression with reduced muscle damage, Gene Ther, 8 (2001) 1264–1270.

[19] European Medicines Agency (EMA), Invega: European Public Assessment Report (EPAR) - Scientific Discussion, 2007. Freely available: https://www.ema.europa.eu/en/documents/scientific-discussion/invega-epar-scientific-discussion_en.pdf.

[20] J.A. Yáñez, C.M. Remsberg, C.L. Sayre, M.L. Forrest, N.M. Davies, Flip-flop pharmacokinetics--delivering a reversal of disposition: challenges and opportunities during drug development, Ther Deliv, 2 (2011) 643–672.

[21] F. Sun, Z. Su, C. Sui, C. Zhang, L. Yuan, Q. Meng, L. Teng, Y. Li, Studies on the acute toxicity, pharmacokinetics and pharmacodynamics of paliperidone derivatives--comparison to paliperidone and risperidone in mice and rats, Basic Clin Pharmacol Toxicol, 107 (2010) 656–662.

[22] U. S. Department of Health and Human Services Food and Drug Administration Center for Drug Evaluation and Research (CDER) Center for Veterinary Medicine (CVM), Bioanalytical Method Validation. Guidance for Industry, 2018. Freely available: https://www.fda.gov/regulatory-information/search-fda-guidance-documents/bioanalytical-method-validation-guidance-industry.

[23] R Core Team, R: A Language and Environment for Statistical Computing. R Foundation for Statistical Computing, in, Vienna, Austria, 2020. Freely available: https://www.r-project.org/.

[24] B.A. Buhren, H. Schrumpf, N.P. Hoff, E. Bölke, S. Hilton, P.A. Gerber, Hyaluronidase: from clinical applications to molecular and cellular mechanisms, Eur J Med Res, 21 (2016) 5.

