## Supplemental Tables and Figures for "Hyaluronidase impacts exposures of long-acting injectable paliperidone palmitate in rodent models"

**SUPPLEMENTARY DATA**

Supplementary Tables

- **Table S1.** PK sampling schedule and number and identity of mice to be sampled following IM and IV dosing.
- **Table S2.** PK sampling schedule and number and identity of rats to be sampled following IM dosing.

Supplementary Figure

- **Figure S1.** Plasma concentration-time PK profile for paliperidone in mice following 3.5 mg/kg IV dose.
- **Figure S2.** Representative hematoxylin and eosin stained sections showing the paliperidone palmitate depot site with muscle or within the adjacent adipose tissue in mouse thighs.

Supplementary Files

- **File S1.** Individual mouse PK data and group PK parameter estimates.
- **File S2.** Individual rat PK data and group PK parameter estimates.

**Table S1. PK sampling schedule and number and identity of mice to be sampled following IM and IV dosing.** For the mouse PK study, timing was relative to the first dosing of M1 in each study group. The schedule of blood sampling and the mouse numbers were applied to each of the four study groups that received IM injections. Mice in the IV injection group were dosed in order from M1 to M16, and blood sampling was done in the order presented in the table to allow for the 6 minute and 15 minute sampling time points.

| Post-dose sampling time point | IM injection, mouse numbers | IV injection, mouse numbers |
|---|---|---|
| 0.1 h (6 min) | --- | M13-M16 |
| 0.25 h (15 min) | M1-M4 | M9-M12 |
| 0.5 h (30 min) | --- | M5-M8 |
| 1 h | M5-M8 | M1-M4 |
| 2 h | M9-M12 | M13-M16 |
| 4 h | M1-M4 | M9-M12 |
| 7 h | M5-M8 | M5-M8 |
| 10 h | M9-M12 | M1-M4 |
| 24 h | M1-M4 | M13-M16 |
| 48 h | M5-M8 | --- |
| 72 h (3 days) | M9-M12 | --- |
| 96 h (4 days) | M1-M4 | --- |
| 168 h (7 days) | M5-M8 | --- |
| 336 h (14 days) | M9-M12 | --- |
| 672 h (28 days) | M1-M4 | --- |

**Table S2. PK sampling schedule and number and identity of rats to be sampled following IM dosing.** For the rat PK study, timing was according to a pre-determined absolute (clock) schedule.

| IM injection groups | | | | | | | | | | | | | | | | |
|---|---|---|---|---|---|---|---|---|---|---|---|---|---|---|---|---|
| **Group** | **Rat number** | **Dose time** | **0.25 h (15 min)** | **1 h** | **2 h** | **4 h** | **7 h** | **10 h** | **24 h** | **48 h** | **72 h (3 days)** | **96 h (4 days)** | **168 h (7 days)** | **360 h (15 days)** | **504 h (21 days)** | **672 h (28 days)** |
| **1** | **1** | **08:00** | 08:15 | 09:00 | 10:00 | 12:00 | 15:00 | 18:00 | 08:00 | 08:00 | 08:00 | 08:00 | 08:00 | 08:00 | 08:00 | 08:00 |
| | **2** | **08:05** | 08:20 | 09:05 | 10:05 | 12:05 | 15:05 | 18:05 | 08:05 | 08:05 | 08:05 | 08:05 | 08:05 | 08:05 | 08:05 | 08:05 |
| | **3** | **08:10** | 08:25 | 09:10 | 10:10 | 12:10 | 15:10 | 18:10 | 08:10 | 08:10 | 08:10 | 08:10 | 08:10 | 08:10 | 08:10 | 08:10 |
| | **4** | **08:15** | 08:30 | 09:15 | 10:15 | 12:15 | 15:15 | 18:15 | 08:15 | 08:15 | 08:15 | 08:15 | 08:15 | 08:15 | 08:15 | 08:15 |
| **2** | **1** | **08:20** | 08:35 | 09:20 | 10:20 | 12:20 | 15:20 | 18:20 | 08:20 | 08:20 | 08:20 | 08:20 | 08:20 | 08:20 | 08:20 | 08:20 |
| | **2** | **08:25** | 08:40 | 09:25 | 10:25 | 12:25 | 15:25 | 18:25 | 08:25 | 08:25 | 08:25 | 08:25 | 08:25 | 08:25 | 08:25 | 08:25 |
| | **3** | **08:30** | 08:45 | 09:30 | 10:30 | 12:30 | 15:30 | 18:30 | 08:30 | 08:30 | 08:30 | 08:30 | 08:30 | 08:30 | 08:30 | 08:30 |
| | **4** | **08:35** | 08:50 | 09:35 | 10:35 | 12:35 | 15:35 | 18:35 | 08:35 | 08:35 | 08:35 | 08:35 | 08:35 | 08:35 | 08:35 | 08:35 |
| **3** | **1** | **08:40** | 08:55 | 09:40 | 10:40 | 12:40 | 15:40 | 18:40 | 08:40 | 08:40 | 08:40 | 08:40 | 08:40 | 08:40 | 08:40 | 08:40 |
| | **2** | **08:45** | 09:00 | 09:45 | 10:45 | 12:45 | 15:45 | 18:45 | 08:45 | 08:45 | 08:45 | 08:45 | 08:45 | 08:45 | 08:45 | 08:45 |
| | **3** | **08:50** | 09:05 | 09:50 | 10:50 | 12:50 | 15:50 | 18:50 | 08:50 | 08:50 | 08:50 | 08:50 | 08:50 | 08:50 | 08:50 | 08:50 |
| | **4** | **08:55** | 09:10 | 09:55 | 10:55 | 12:55 | 15:55 | 18:55 | 08:55 | 08:55 | 08:55 | 08:55 | 08:55 | 08:55 | 08:55 | 08:55 |
| | **4** | **11:15** | 11:21 | 11:30 | 11:45 | 12:15 | 13:15 | 15:15 | 18:15 | 21:15 | 11:15 | | | | | |

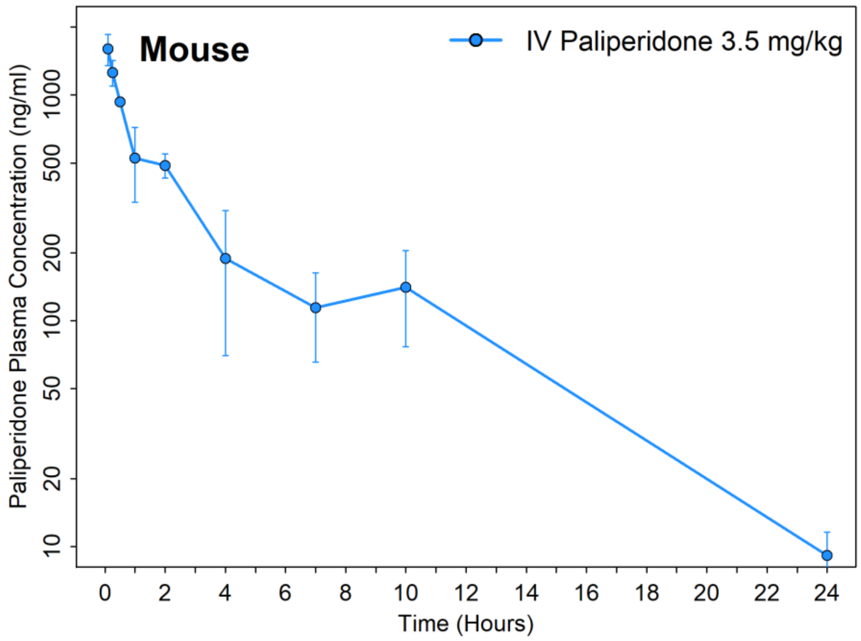


**Figure S1. Plasma concentration-time PK profile for paliperidone in mice following a 3.5 mg/kg dose administered by IV injection.** Data point represent mean values (n = 3 samples per time) and error bars represent standard deviation.

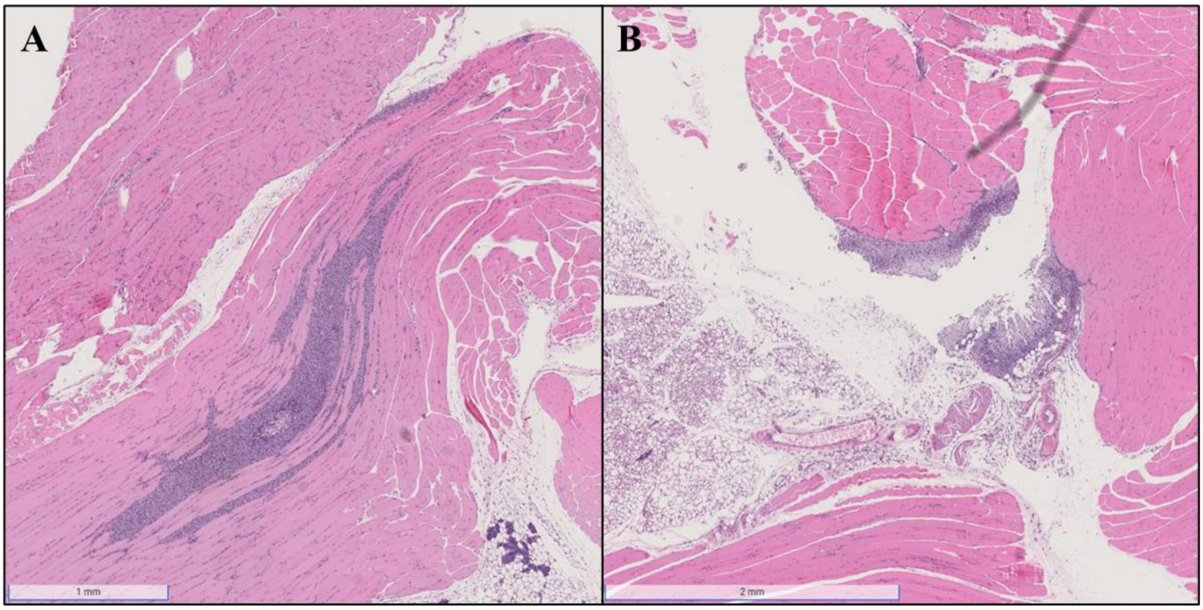


**Figure S2. Representative hematoxylin and eosin stained sections showing the paliperidone palmitate depot site with muscle (A) or within the adjacent adipose tissue (B) in mouse thighs.** Panel A shows the depot site in a mouse thigh that was injected with paliperidone palmitate with buffer only (no hyaluronidase), 336 hours (2 weeks) post injection. The depot site, visible by the mass of macrophages, is clearly within the muscle tissue. Panel B shows the depot site in a mouse thigh that was also injected with paliperidone palmitate with buffer only (no hyaluronidase), 168 hours (1 week) post injection. Here, the paliperidone palmitate was either directly injected into the adipose tissue or possibly displaced from the muscle tissue, for example during muscle contraction.
